# Rare protein-altering variants in *ANGPTL7* lower intraocular pressure and protect against glaucoma

**DOI:** 10.1101/677443

**Authors:** Yosuke Tanigawa, Michael Wainberg, Juha Karjalainen, Tuomo Kiiskinen, Susanna Lemmelä, Joni A. Turunen, Robert Graham, Aki S. Havulinna, Markus Perola, Aarno Palotie, FinnGen, Mark J. Daly, Manuel A. Rivas

## Abstract

Protein-altering variants that are protective against human disease provide *in vivo* validation of therapeutic targets. Here we use genotyping data in UK Biobank and FinnGen to conduct a search for protein-altering variants conferring lower intraocular pressure (IOP) and protection against glaucoma. Through protein-altering variant association analysis we find a missense variant in UK Biobank (rs28991009 (MAF=0.8%) genotyped in 81,527 individuals with measured IOP and an independent set of 4,269 glaucoma patients and 251,355 controls) that significantly lowers IOP (β = −0.73 mmHg for heterozygotes, −2.96 mmHg for homozygotes, *P* = 1 × 10^−13^) and is associated with 34% reduced risk of glaucoma (*P* = 0.005). In FinnGen, we identify an *ANGPTL7* missense variant at a greater than 50-fold increased frequency in Finland compared with other populations (rs147660927, p.Arg220Cys, MAF Finland = 4.1%), which was genotyped in 5,177 glaucoma patients and 130,461 controls and is associated with 30% lower glaucoma risk (*P* = 1 × 10^−9^). We further find three rarer variants in UK Biobank, including a protein-truncating variant, which confer a strong composite lowering of IOP (*P* = 0.002), suggesting the protective mechanism likely resides in the loss of an interaction or function. Our results support inhibition or down-regulation of ANGPTL7 as a therapeutic strategy for glaucoma

## Introduction

Intraocular pressure is currently the sole modifiable risk factor and predictive measure for glaucoma^1,2,3,4^ (**Supplementary Figure S1**).Genome-wide association studies(GWAS) have commonly used genetic associations to this endophenotype that exhibits high genetic correlation (rg = 0.71) to glaucoma, as an approach to prioritize genetic variants likely to contribute to disease risk^5^. A total of 53 independent loci have been unequivocally implicated in glaucoma^5^. For these discoveries, like most GWAS results, it has proven challenging to infer the functional consequences of common variant associations beyond cases where protein-altering variants have been directly implicated. Protein-altering variants, generally the strongest-acting genetic variants in medical genetics, include nonsynonymous substitutions and protein-truncating variants, and understanding their functional consequences provides insight into the therapeutic effects of inhibiting or down-regulating the gene in which they reside^6^. Thus, identifying protein-altering variants that confer protection from disease holds particular promise for identifying therapeutic targets.

Here we leverage two cohorts that provide complementarity for glaucoma gene discovery. First, UK Biobank has obtained intraocular pressure (IOP) measurements in approximately 128,000 individuals in addition to case-control status for glaucoma from hospital in-patient and verbal questionnaire data in over 500,000 individuals^7, 8^. Second, FinnGen has directly genotyped and aggregated disease outcomes in over 135,000 individuals from Finland, an isolated population with recent bottlenecks that offers an unprecedented advantage for studying rare variants in complex diseases^9^. With clinic-based recruitment focused on several areas including ophthalmology, and with 32.4% of the collection above age 70, FinnGen is particularly well-powered for aging-associated endpoints. We therefore conduct targeted association analysis with IOP in UK Biobank (N = 81,534) to identify protein-altering variants that reduce IOP, and test whether those variants, or others in the same genes, also confer protection to glaucoma in FinnGen (5,177 cases and 130,461 controls) and UK Biobank (4,269 cases and 251,355 controls not included in the IOP sample). Analysis of an allelic series of protein-altering variants in *ANGPTL7* in 9,446 glaucoma patients and over 350,000 controls identifies a significant lowering effect on IOP and protective association with glaucoma. By analyzing putative loss-of-function variants, we find concordant effect directions with the nonsynonymous substitutions, suggesting that the protective mechanism may reside in the loss of gene function.

## Results

### Protein-altering variant association analysis

We conducted protein-altering variant association analysis with IOP, as measured via Goldmann-correlated tonometry, in 81,527 British individuals in UK Biobank dataset (Methods)^10^. Across 41,632 rare (0.01 % < MAF < 1%) protein-altering variants outside of the MHC region with genotyping array data in UK Biobank, we used a generalized linear model implemented in PLINK^11^ to scan for variants with IOP-lowering effects. We identified one protein-altering variant significantly associated with lower IOP below a Bonferroni-corrected *P* value < 1.0×10^−6^, a nonsynonymous substitution (p.Gln175His) in *ANGPTL7* (*P* = 1.47×10^−13^, β = −0.200 SD 95% CI: [−0.253, −0.147], −0.73 mHg for heterozygotes, −2.96 mmHg for homozygotes, Table 1,2, Supplementary Figure S2).

**Table 1.** *ANGPTL7* IOP protein-altering variant association in UK Biobank. . Variant includes chromosome, position, reference, and alternate allele (hg19). rsID - the rs identifier of the genetic variant. HGVSp - the HGVS protein sequence name. MAF - the minor allele frequency in UK Biobank British population. Beta - estimated regression coefficient with 95% confidence intervals. *P* - p-value of association.

| Variant | rsID | HGVSp | MAF | Beta SD [95% CI] | $P$ |
| --- | --- | --- | --- | --- | --- |
| 1:11252357:A:G | rs200058074 | p.Gln136Arg | .054 % | -.038 [-.26, .18] | $7.3 \times 10^{-1}$ |
| 1:11252369:G:A | rs28991002 | p.Arg140His | .25 % | -.16 [-.25, -.061] | $1.3 \times 10^{-3}$ |
| 1:11253684:G:T | rs28991009 | p.Gln175His | .81 % | -.20 [-.25, -.15] | $1.4 \times 10^{-13}$ |
| 1:11253688:C:T | rs143435072 | p.Arg177Ter | .041 % | -.26 [-.51, -.014] | $3.8 \times 10^{-2}$ |

**Table 2.** ANGPTL7 allelic series summary. . Variant: HGVSp amino acid nomenclature. Dosage - genotype of individuals. rsID - rs identifier. Carrier frequency in UK - carrier frequency in UK Biobank for the respective genotype dosage. Carrier frequency in Finland - carrier frequency in FinnGen for the respective genotype dosage. N with IOP (UK Biobank) - number of individuals in UK Biobank with intraocular pressure measurements corresponding to the genotype dosage. IOP (mmHg) [95% CI] - unstandardized estimates of effect size on intraocular pressure measurements (NB: standardized estimated effect sizes may have lower p-values due to normalization procedure). OR for glaucoma - estimate odds ratio on glaucoma risk for the respective genotype dosage. NS non-significant (p > 0.1).

| Variant | Dosage | rsID | Carrier frequency in UK | Carrier frequency in Finland | N with IOP (UK Biobank) | IOP (mmHg) [95% CI] | OR for glaucoma |
| --- | --- | --- | --- | --- | --- | --- | --- |
| Gln175His | Gln/His | rs28991009 | 1.43% | 0.24% | 1355 | -0.73 [-0.95, -0.52] *** | 0.66 ** |
| Gln175His | His/His |  | 0.01% | NA | 5 | -2.95 [-6.43, 0.530] . | NS |
| Arg177Ter | Arg/Ter | rs143435072 | 0.07% | NA | 62 | -0.94 [-1.93, 0.0445] . | NS |
| Arg140His | Arg/His | rs28991002 | 0.51% | 0.35% | 427 | -0.48 [-0.86, -0.10] * | NS |
| Gln136Arg | Gln/Arg | rs200058074 | 0.11% | NA | 80 | -0.21 [-1.08, 0.66] | NS |
| Arg220Cys | Arg/Cys | rs147660927 | NA | 7.82% | NA | NA | 0.72 *** |
| Arg220Cys | Cys/Cys |  | NA | 0.21% | NA | NA | 0.47 . |

Significant codes: 0 '\*\*\*' 0.001 '\*\*' 0.01 '\*' 0.05 '.' 0.1 ' ' 1

Based on this finding, we assessed whether any additional rare variant associations in *ANGPTL7* were present. We found three additional independent rare protein-altering variants in *ANGPTL7* (MAF < 0.25%) including a premature stop-gain allele (p.Arg177Ter). Collectively, these three variants showed a marginally significant association with lower IOP (*P* = 0.0023, Table 1), with the stop-gain allele p.Arg177Ter also showing a marginally significant effect on its own (*P* = 0.038). Genotyping intensity plots and the concordance of genotype calls from array and whole-exome sequencing data were manually inspected to ensure high quality and consistent genotyping (Supplementary Figure S3, Supplementary Table S1, Methods) and alleles were confirmed to be independent (pairwise r^2^ < 10^−4^ for each, Supplementary Table S2, Methods). Using inverse variance weighted meta-analysis method, the combined effect of those four variants are estimated to be −0.19 ([95% CI: −0.23 - −0.14] SD, *P* = 3.4×10^−16^; −0.67 mmHg [95% CI: −0.85 - −0.49], Supplementary Figure S4, Supplementary Tables S3-4, Methods).

Given these findings, we next asked whether any of these putative IOP-lowering genetic variants showed effects consistent with reducing glaucoma risk in an independent set of British individuals that do not have IOP measures (4,269 cases and 251,355 controls). For p.Gln175His in *ANGPTL7*, we estimated that the variant lowers glaucoma risk by 34% (*P* = 0.00543; OR = 0.66 [95% CI: 0.366 - 0.954]). The three additional protein-altering variants did not significantly confer protection against glaucoma (burden test *P* = 0.55), although we expect this to be a result of limited power in the binary case-control setting.

### Independent Finnish-enriched missense allele in *ANGPTL7*

We then sought evidence from the FinnGen dataset that either the same or novel Finnish-enriched protein-altering variants would confirm the association of *ANGPTL7* with protection from glaucoma. This additional *ANGPTL7* association data, obtained in 5,177 glaucoma patients and 130,461 controls, provided strong support that protein-altering variants in *ANGPTL7* protect against glaucoma (case definitions described in Supplementary Table S5). Specifically, we found that the p.Gln175His substitution has nominal evidence of association (*P* = 0.031, OR = 0.49) despite the variant only being present at a minor allele frequency of 0.1% in this Finnish cohort (8-fold depleted compared to UK Biobank). The remaining protein-altering variants in *ANGPTL7* tested in UK Biobank were not found in the FinnGen dataset. Confirmation of an *ANGPTL7* effect on glaucoma risk was seen in data from an independent Finnish-specific protein-altering nonsynonymous substitution, p.Arg220Cys, which was strongly associated with protection from glaucoma (*P* = 1.0×10^−9^, OR = 0.72 [95% CI: 0.64 - 0.80], Supplementary Figure S5). Of note, this observation is advantaged by the property that p.Arg220Cys is found at a greater than 50-fold increased frequency in Finland compared with other populations^12^, reinforcing the value of isolated, bottlenecked populations in which the allele frequency spectrum is intensely concentrated on the minority of variants passing through the bottleneck.

While registry-based diagnoses in FinnGen do not yet contain detailed ophthalmologic records, a subset of 2695 glaucoma patients had been recorded as having primary open-angle glaucoma (POAG). In this sub-group a stronger effect was observed (OR=0.68) versus those glaucoma cases without a definitive record of POAG (OR=0.77 [95% CI: 0.67 - 0.88]), reminiscent of the stronger risk effects seen at the myocilin (*MYOC*) gene and other established genes in the POAG subgroup. Given the Finnish enrichment of the known strong glaucoma risk allele, p.Gln368Ter, in *MYOC* (MAF in Finland = 0.3%, MAF in Non-Finnish European = 0.16%, reference sequence: NM_00026), we next asked whether carriers have risk reduced if they carry *ANGPTL7* p.Arg220Cys. In FinnGen, we estimate that 7.7% of individuals carriers for *MYOC* p.Gln368Ter variant are POAG cases in comparison to 2% for non-carriers. In the presence of *ANGPTL7* p.Arg220Cys, only 1.3% of individuals are POAG cases, and only 1 of 70 (1.4%) who carry both *MYOC* risk and *ANGPTL7* protective variants were POAG cases (Supplementary Table S6). This suggests *ANGPTL7* protection extends to the *MYOC* risk group but the small counts preclude any definitive statement regarding interaction - given the limited number of double-carriers, larger case-control series are needed to refine our understanding as to whether *ANGPTL7* p.Arg220Cys variant modifies the glaucoma risk conferred by p.Gln368Ter in *MYOC*.

Access to genotype data in over 300,000 individuals from the UK and 135,000 from the Finnish group enabled us to identify rare protein-altering homozygotes. In UK Biobank we found 28 individuals homozygous for the 175His allele, consistent with Hardy-Weinberg expectation (n=22.6), where we estimated a 2.96 mm Hg drop in IOP compared to the mean IOP levels. Furthermore, the oldest reached the age of 80 and one of the 28 died (age 65). In FinnGen we found 268 individuals homozygous for the 220C allele, the oldest reached the age of 98, with no depletion of homozygotes compared with Hardy-Weinberg equilibrium expectation. There was no significant association of the homozygous genotype with a decreased lifespan. Together this indicates that having two copies with 175His or 220Cys in *ANGPTL7* is compatible with normal lifespan. To assess the potential impacts of those protein-truncating variants on reproductive fitness we assessed the association of p.Gln175His with the number of live births and number of children fathered and found no significant association (*P* > 0.05/4, Supplementary Table S7). Hence, we did not find any severe medical consequences that would be of obvious concern in developing a therapeutic to mimic the effect of these alleles.

The combined significance of *ANGPTL7* protein-altering variants across more than 435,000 samples tested is *P* = 3.4×10^−16^ for IOP and *P* = 1.9×10^−11^ for glaucoma using inverse-variance weighted method (Supplementary Figure S6).

### Transcript and protein expression

*ANGPTL7*, a five-exon protein-coding gene, encodes the Angiopoietin-related protein 7, which is expressed in several human tissues including the trabecular meshwork, cornea and retina ^13–15^. Recently, *ANGPTL7* overexpression in primary human trabecular meshwork cells was found to alter the expression of relevant trabecular meshwork proteins of the extracellular matrix, including fibronectin, collagens type I, IV, and V, myocilin, versican, and MMP1, and ANGPTL7 protein was increased as the disease progressed in POAG beagle dogs^13^. We examined proteomics expression data in normal tissues and cell lines from ProteomicsDB and MOPED^16, 17^ and found vitreous humor tissue-specific expression of ANGPTL7 (log10 ppm = 1, Supplementary Figure S7). These data suggest that the eye is the relevant tissue type in understanding the functional consequences of the variants discovered in this study. Further work is dissecting the role of *ANGPL7* in all possible cell types in the eye is warranted.

## Discussion

This study establishes strong genetic evidence for the involvement of *ANGPTL7* in glaucoma risk in which a powerful allelic series, including all low-frequency nonsynonymous substitutions and a single premature stop-gain substitution, is conclusively associated with reduced disease risk and endophenotype-lowering effects. Increasing evidence is associating ANGPTL proteins to cardiometabolic phenotypes^18–22^. Although it has been recently proposed that ANGPTL7 levels are increased in obesity and reduced after physical exercise, we do not observe any evidence of genetic association in either UK Biobank or FinnGen to support this hypothesis^23^. In the context of the other established variants in glaucoma, including the protein-truncating variants in *MYOC*, p.Gln175His and the 57-fold Finnish enriched p.Arg220Cys variant in *ANGPTL7* exert a comparable protective effect. Because of the strong protective effect associated with the *ANGPTL7* coding variants, studies of ANGPTL7 inhibition and the specific action of these variant proteins should be useful in understanding the mechanism by which protection to glaucoma disease occurs and whether this reveals a promising therapeutic opportunity similar to that which has been realized from the examples of PCSK9, *APOC3* and cardiovascular disease^24–26^. The phenotype and longevity profile of *ANGPTL7* homozygotes suggest that this is likely to be a safe strategy for therapeutic intervention.

## Acknowledgments

This research has been conducted using the UK Biobank Resource under Application Number 24983, “Generating effective therapeutic hypotheses from genomic and hospital linkage data” (http://www.ukbiobank.ac.uk/wp-content/uploads/2017/06/24983-Dr-Manuel-Rivas.pdf). Based on the information provided in Protocol 44532 the Stanford IRB has determined that the research does not involve human subjects as defined in 45 CFR 46.102(f) or 21 CFR 50.3(g). All participants of UK Biobank provided written informed consent (more information is available at https://www.ukbiobank.ac.uk/2018/02/gdpr/). For the Finnish Institute of Health and Welfare (THL) driven FinnGen preparatory project (here called FinnGen), all patients and control subjects had provided informed consent for biobank research, based on the Finnish Biobank Act. Alternatively, older cohorts were based on study specific consents and later transferred to the THL Biobank after approval by Valvira, the National Supervisory Authority for Welfare and Health. Recruitment protocols followed the biobank protocols approved by Valvira. The Ethical Review Board of the Hospital District of Helsinki and Uusimaa approved the FinnGen study protocol Nr HUS/990/2017. The FinnGen preparatory project is approved by THL, approval numbers THL/2031/6.02.00/2017, amendments THL/341/6.02.00/2018, THL/2222/6.02.00/2018 and THL/283/6.02.00/2019. All DNA samples and data in this study were pseudonymized. We thank all the participants in the UK Biobank and Finnish Biobanks used in FinnGen studies. This work was supported by National Human Genome Research Institute (NHGRI) of the National Institutes of Health (NIH) under awards R01HG010140 (M.A.R.). The content is solely the responsibility of the authors and does not necessarily represent the official views of the National Institutes of Health. We thank Sirpa Soini, Fatima Rodriguez and David Amar for invaluable feedback on the manuscript. Y.T. is supported by Funai Overseas Scholarship from Funai Foundation for Information Technology and the Stanford University School of Medicine. FinnGen is supported by Abbvie, Astra Zeneca, Biogen, Celgene, Genentech, GSK, Merck, Pfizer, and Sanofi. M.A.R. is supported by Stanford University and a National Institute of Health center for Multi- and Trans-ethnic Mapping of Mendelian and Complex Diseases grant (5U01 HG009080).

## Author information

### Author contributions

M.A.R. and M.J.D. conceived and designed the study. M.P. designed the FinnGen preparatory project in collaboration with A.P and M.J.D. M.A.R., Y.T., J.K., and M.J.D. carried out the statistical and computational analyses. M.A.R., Y.T., J.K., T.K., J.A.T., and M.J.D. carried out quality control of the data. The manuscript was written by M.A.R., Y.T., and M.J.D.; and revised by all the co-authors. All co-authors have approved of the final version of the manuscript.

### Data availability

Data is displayed in the Global Biobank Engine (https://biobankengine.stanford.edu). Analysis scripts and notebooks are available on GitHub at https://github.com/rivas-lab/ANGPTL7.

## Methods

### Genome-wide association analysis in UK Biobank

For British individuals (n = 337,151) in UK Biobank as described elsewhere^8^, we applied genome-wide association analysis for directly genotyped variants were applied to Goldmann-correlated intraocular pressure described more detailed earlier^10^ (right, UK Biobank Field ID 5255, Global Biobank Engine phenotype ID: INI5255) and glaucoma (Global Biobank Engine phenotype ID: HC276), which is previously defined as a part of “high confidence” disease outcome phenotypes by combining disease diagnoses from the UK National Health Service Hospital Episode Statistics with self-reported diagnoses questionnaire^8^), using generalized linear model association analysis implemented in PLINK v2.00aLM (2 April 2019) with age, sex, types of genotyping array, and the first 4 genotype principal components as described elsewhere^11, 27^. The glaucoma phenotype was previously defined as a part of “high confidence” disease outcome phenotypes by combining disease diagnoses (UK Biobank Field ID 41202) from the UK National Health Service Hospital Episode Statistics (ICD10 codes: H40.[0-6,8,9], H42.8, and Q15.0) with self-reported non-cancer diagnosis questionnaire (UK Biobank Field ID 20002)^8^.

### Genotyping quality control in UK Biobank

#### Manual inspection of intensity plots

For the identified rare (0.01 % < MAF < 1%) protein-altering variants in *ANGPTL7* (reference sequence: NM_021146), we generated and inspected intensity plots with McCarthy Group’s ScatterShot using “UKB - All Participants” module^28^.

#### Variant-calling consistency analysis

For individuals with whole-exome sequencing data (n = 49,960), we extracted the genotype calls of coding variants in *ANGPTL7* using PLINK v2.00aLM (2 April 2019) and compared the consistency between array-genotyped dataset and whole-exome sequencing dataset^11, 29^.

### Independence analysis of alleles

#### Pairwise r^2^ computation within British individuals in UK Biobank

We computed pairwise r^2^ for the identified rare protein-altering variants in *ANGPTL7* within British individuals in UK Biobank using PLINK v1.90b6.7 64-bit (2 Dec 2018) with --ld <variant_ID_1> <variant_ID_1> hwe-midp subcommand^11^.

#### Number of individuals with the combination of genotypes in UK Biobank

Using the extracted genotype calls from for the identified rare protein-altering variants in *ANGPTL7* (see Variant-calling consistency analysis section), we counted the number of British individuals by the combination of genotypes. We computed the expected number of individuals under Hardy-Weinberg equilibrium model and the independence assumption:

- The expected frequencies of REF/REF, REF/ALT, and ALT/ALT carriers are (1 - AF)^2^, 2 × AF(1-AF), and AF^2^, respectively.
- The expected genotyping rate is independently estimated by the observed genotyping rate for each variant.
- The expected frequency of the combination of genotypes is computed under the independent assumption among alleles

### Combined effect size estimates of rare protein-altering variants

Using the inverse-variance method for fixed-effects model implemented in R meta package, we performed a meta-analysis of the estimated effect sizes (BETAs) of protein-altering variants on intraocular pressure and glaucoma. For glaucoma, the results are reported as odds ratio. 95% confidence intervals are reported for IOP and glaucoma.

### Association analysis with reproductive fitness

Using the number of live births (UK Biobank Field ID: 2734, Global Biobank Engine phenotype ID: INI2734) and the number of children fathered (UK Biobank Field ID: 2405, Global Biobank Engine phenotype ID: INI2405), we performed association analysis for the four identified protein-altering variants using R script with age, types of genotyping array, and the first 4 genotype principal components as covariates.

### Homozygous carrier analysis

For UK Biobank British individuals, we extracted the genotype calls with PLINK v2.00aLM (2 April 2019) and identified homozygous carrier of 175H allele^11^. We examined the year of birth (UK Biobank Field ID 34) and age at death (UK Biobank Field ID 40007)^7^.

## Supplementary Tables and Figures

**Supplementary Figure S1.**
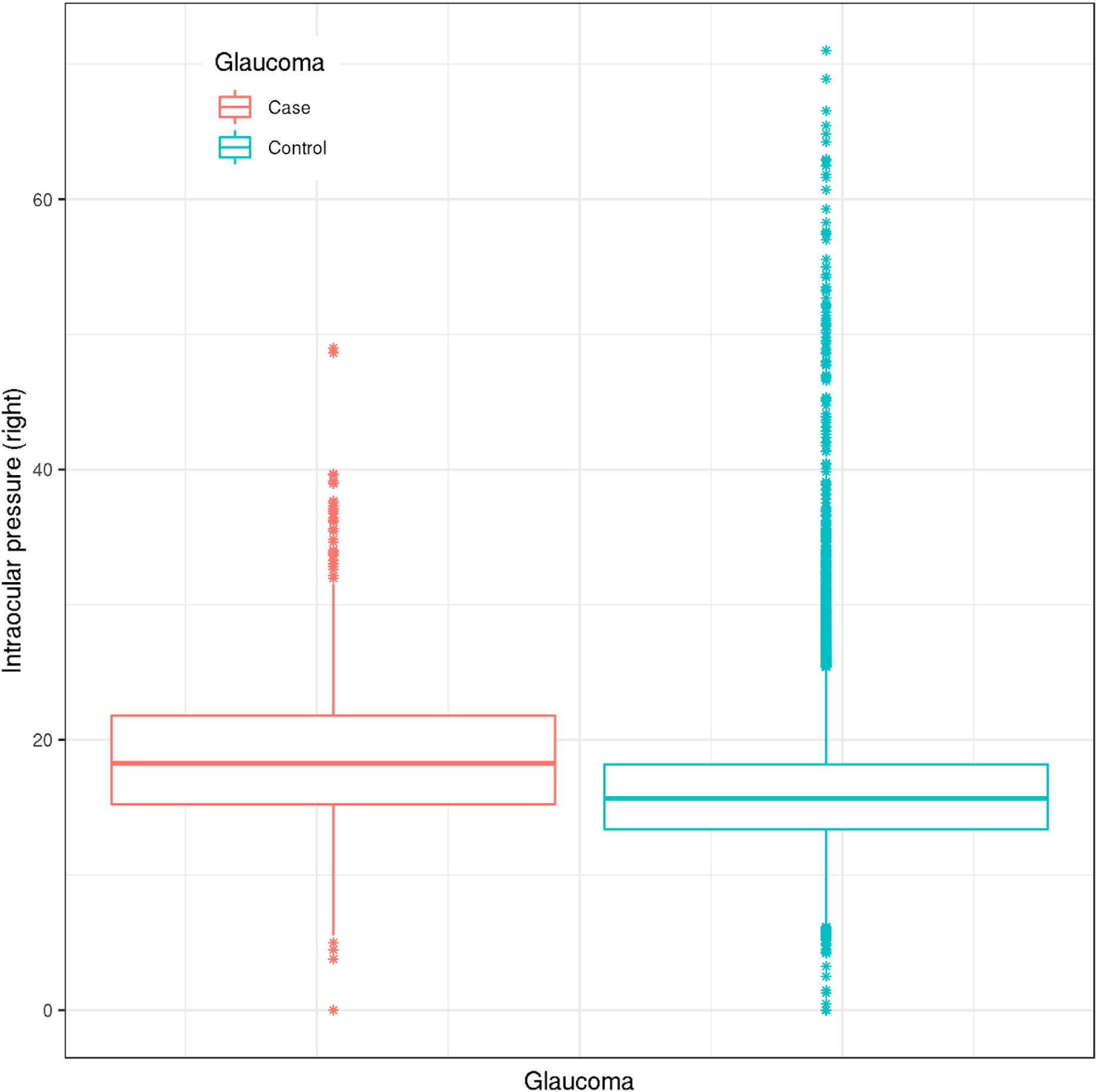
Phenotype distribution of Intraocular pressure stratified by glaucoma disease status displayed as a Tukey’s box plot. The middle bold horizontal line represents the median, the lower and upper hinges show the first and third quartiles, the lower and upper whiskers represent 1.5 * interquartile range from the hinges. The data points beyond whiskers are plotted individually.

**Supplementary Figure S2.**
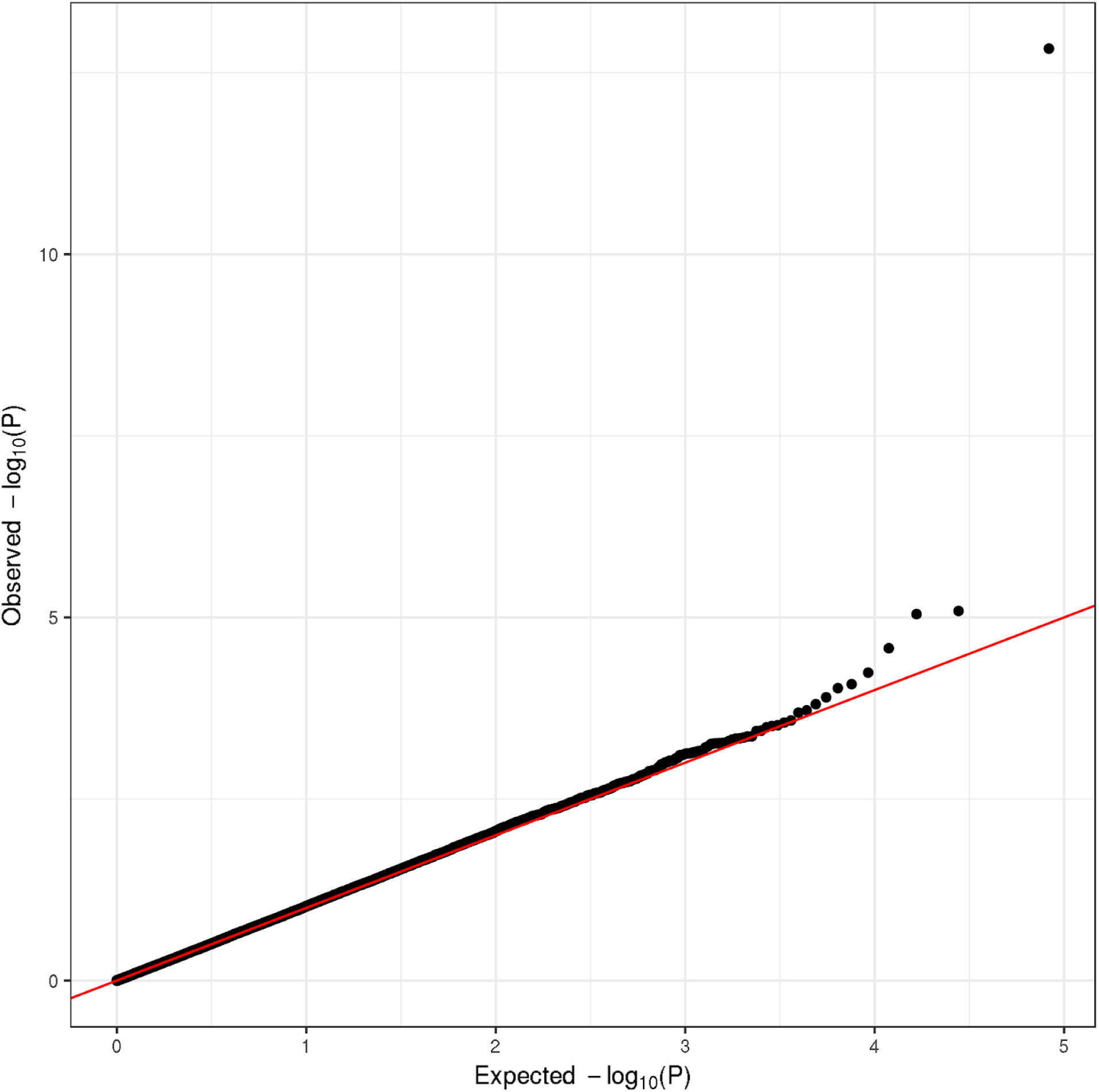
The protein-altering variant GWAS QQ plot for intraocular pressure. The variants outside of MHC region with 0.01 % < MAF < 1% are included in the analysis.

**Supplementary Figure S3.**
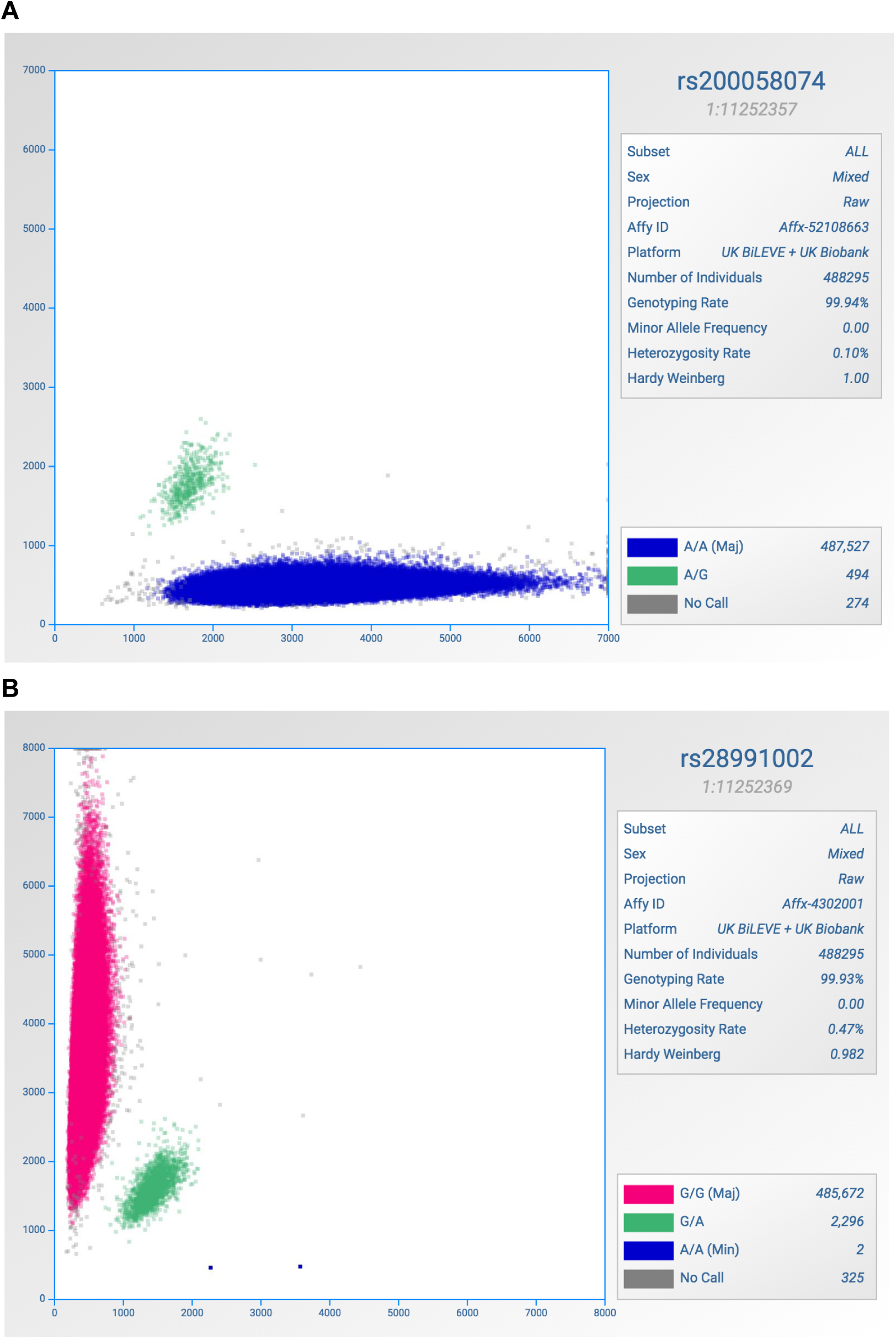
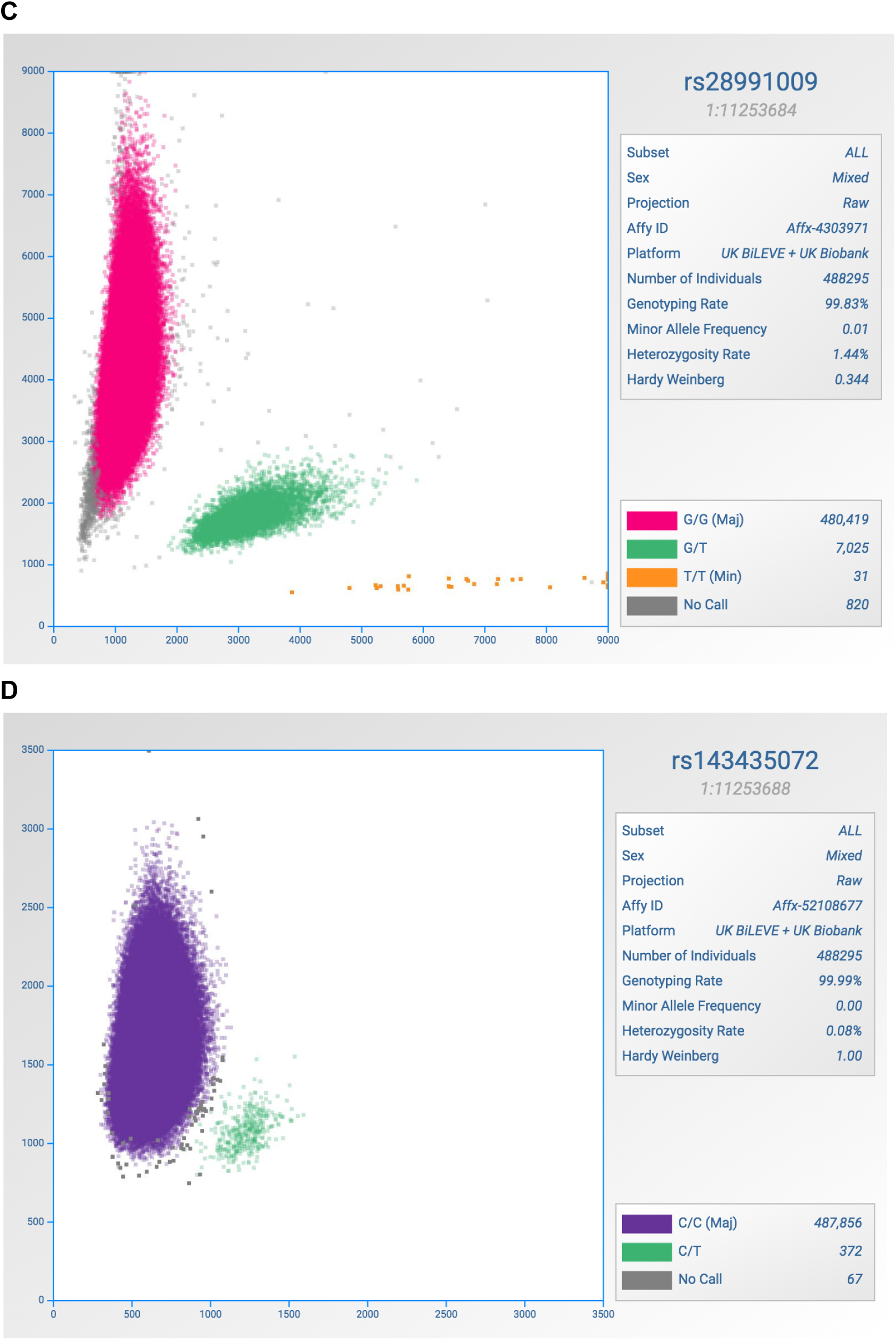
The intensity plots for ANGPTL7 protein-altering variants with 0.01 % < MAF < 1%. **(A)** rs200058074 (p.Gln136Arg). **(B)** rs28991002 (p.Arg140His). **(C)** rs28991009 (p.Gln175His). **(D)** rs143435072 (p.Arg177Ter)

**Supplementary Table S1.** Consistency of the genotype call for four protein-altering variants in *ANGPTL7* between genotyping array and exome sequencing data. Variant including chromosome, position, reference, and alternate allele (hg19), the rs identifier of the genetic variant (rsID), consequence of the variation (Consequence), genotype call from array (Array) and exome data (Exome), and the number of individuals (N). Inconsistent variant calls are highlighted in bold font.

| Variant | rsID | Consequence | Array | Exome | N |
| --- | --- | --- | --- | --- | --- |
| 1:11252357:A:G | rs200058074 | p.Gln136Arg | A/G | A/G | 42 |
| 1:11252357:A:G | rs200058074 | p.Gln136Arg | A/A | A/A | 49837 |
| 1:11252357:A:G | rs200058074 | p.Gln136Arg | NA | A/G | 1 |
| 1:11252357:A:G | rs200058074 | p.Gln136Arg | NA | A/A | 29 |
| 1:11252369:G:A | rs28991002 | p.Arg140His | A/A | A/A | 1 |
| 1:11252369:G:A | rs28991002 | p.Arg140His | G/A | G/A | 218 |
| 1:11252369:G:A | rs28991002 | p.Arg140His | **G/A** | **G/G** | **1** |
| 1:11252369:G:A | rs28991002 | p.Arg140His | G/G | G/G | 49,663 |
| 1:11252369:G:A | rs28991002 | p.Arg140His | NA | G/G | 26 |
| 1:11253684:G:T | rs28991009 | p.Gln175His | T/T | T/T | 4 |
| 1:11253684:G:T | rs28991009 | p.Gln175His | G/T | G/T | 720 |
| 1:11253684:G:T | rs28991009 | p.Gln175His | G/G | G/G | 49,093 |
| 1:11253684:G:T | rs28991009 | p.Gln175His | NA | G/T | 1 |
| 1:11253684:G:T | rs28991009 | p.Gln175His | NA | G/G | 91 |
| 1:11253688:C:T | rs143435072 | p.Arg177Ter | C/T | C/T | 41 |
| 1:11253688:C:T | rs143435072 | p.Arg177Ter | **C/T** | **C/C** | **1** |
| 1:11253688:C:T | rs143435072 | p.Arg177Ter | C/C | C/C | 49,857 |
| 1:11253688:C:T | rs143435072 | p.Arg177Ter | NA | C/C | 10 |

**Supplementary Table S2.** Number of individuals stratified by genotype of rare (0.01% < MAF < 1%) protein-altering variants in *ANGPTL7*. The combination of genotypes is shown in the first four columns (rs200058074, rs28991002, rs28991009, and rs143435072) as well as the number of British individuals with the genotype combination in UK Biobank (n_observed). The expected number of individuals are computed under Hardy-Weinberg equilibrium model and the independence assumption (n_expected, Method).

| rs200058074 | rs28991002 | rs28991009 | rs143435072 | n_observed | n_expected |
| --- | --- | --- | --- | --- | --- |
| A/A | G/G | G/G | C/C | 328,492.00 | 329,401.21 |
| A/A | G/G | G/T | C/C | 5,395.00 | 5,390.72 |
| A/A | G/A | G/G | C/C | 1,693.00 | 1,672.69 |
| A/A | G/G | NA | C/C | 471.00 | 470.71 |
| A/G | G/G | G/G | C/C | 356.00 | 353.03 |
| A/A | G/G | G/G | C/T | 270.00 | 267.80 |
| A/A | NA | G/G | C/C | 194.00 | 192.47 |
| NA | G/G | G/G | C/C | 192.00 | 188.77 |
| A/A | G/G | G/G | NA | 37.00 | 38.13 |
| A/A | G/G | T/T | C/C | 28.00 | 22.06 |
| A/A | G/A | G/T | C/C | 8.00 | 27.37 |
| A/G | G/G | G/T | C/C | 4.00 | 5.78 |
| A/A | NA | G/T | C/C | 2.00 | 3.15 |
| A/A | G/A | G/G | NA | 2.00 | 0.19 |
| A/A | G/G | NA | C/T | 2.00 | 0.38 |
| NA | G/A | G/G | C/C | 1.00 | 0.96 |
| A/A | A/A | G/G | C/C | 1.00 | 2.12 |
| A/A | G/A | G/G | C/T | 1.00 | 1.36 |
| A/A | G/G | G/T | C/T | 1.00 | 4.38 |
| A/G | G/G | NA | C/C | 1.00 | 0.50 |

**Supplementary Figure S4:**
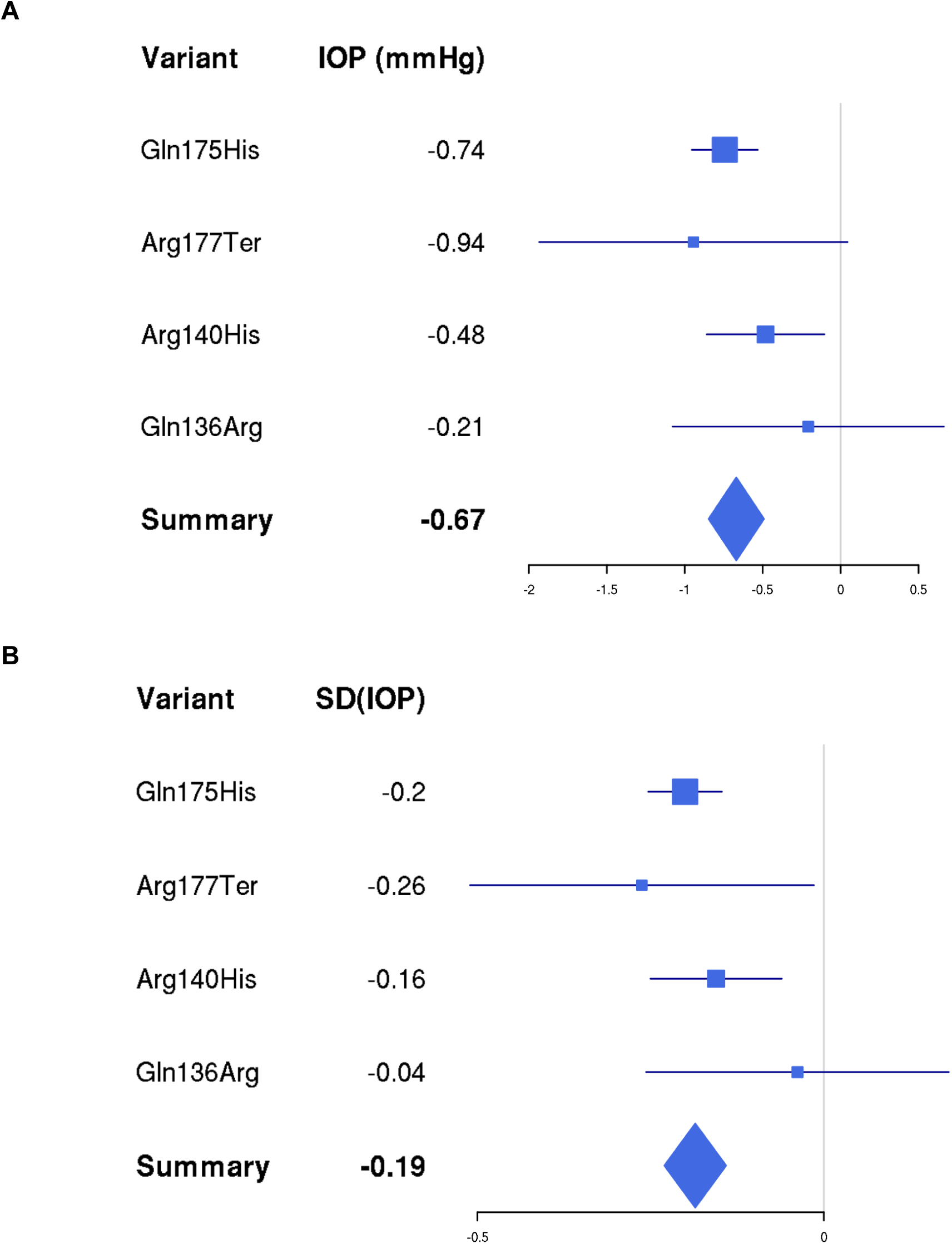
The combined effect size estimates of rare protein-altering variants. The estimates of effect size on unstandardized intraocular pressure measurements (IOP [mmHg]) and normalized intraocular pressure measurements (SD[IOP]) are shown for the four protein-altering variants in *ANGPTL7*. The combined effect size estimate is summarized using inverse-variance method under the fixed effect model (Methods).

**Supplementary Table S3.** The combined effect size estimates of rare protein-altering variants. The estimates of effect size on normalized intraocular pressure measurements (SD [IOP]) and their 95% confidence interval (95% CI) are shown for the four protein-altering variants in *ANGPTL7*. The combined effect size estimate is summarized using inverse-variance method under the fixed effect model (Methods). %W (fixed) column indicates the relative weights used in the fixed effect model. The bottom right cell indicates the P-value of combined effect size model.

| Variant | SD (IOP) | 95% CI | %W(fixed) |
| --- | --- | --- | --- |
| Gln175His | -0.2003 | [-0.2534; -0.1471] | 70.5 |
| Arg177Ter | -0.2626 | [-0.5107; -0.0144] | 3.2 |
| Arg140His | -0.1555 | [-0.2502; -0.0607] | 22.1 |
| Gln136Arg | -0.0380 | [-0.2564; 0.1804] | 4.2 |
| **Fixed effect model** | -0.1856 | [-0.2302; -0.1410] | P-value: $3.4 \times 10^{-16}$ |

**Supplementary Table S4.** The combined effect size estimates of rare protein-altering variants. The estimates of effect size on unstandardized intraocular pressure measurements (IOP [mmHg]) and their 95% confidence interval (95% CI) are shown for the four protein-altering variants in *ANGPTL7*. The combined effect size estimate is summarized using inverse-variance method under the fixed effect model (Methods). %W (fixed) column indicates the relative weights used in the fixed effect model. The bottom right cell indicates the P-value of combined effect size model.

| Variant | IOP (mmHg) | 95% CI | %W(fixed) |
| --- | --- | --- | --- |
| Gln175His | -0.7430 | [-0.9547; -0.5314] | 70.5 |
| Arg177Ter | -0.9444 | [-1.9334; 0.0445] | 3.2 |
| Arg140His | -0.4816 | [-0.8593; -0.1039] | 22.1 |
| Gln136Arg | -0.2078 | [-1.0785; 0.6630] | 4.2 |
| **Fixed effect model** | -0.6694 | [-0.8471; -0.4917] | P-value: $1.6 \times 10^{-13}$ |

**Supplementary Table S5:** Glaucoma definitions in FinnGen. ICD-codes are used in the Finnish hospital discharge and cause-of-death registries. ATC-codes are used in the Social Insurance Institution prescription drug purchase registry. All endpoint definitions in the FinnGen phenome-wide association analysis are available online (https://www.finngen.fi/fi/node/68).

| Phenotype name | ICD-10 | ICD-9 | ICD-8 | ICD-8<br>exclude | ATC (drug<br>purchases) |
| --- | --- | --- | --- | --- | --- |
| Glaucoma (broad definition, combination of all the subphenotypes below) | Any of the below | Any of the below | Any of the below |  | S01EA |
| Glaucoma suspect | H40.0 | 365.0 | 375.97 |  |  |
| Primary open-angle glaucoma | H40.1 | 365.1 | 375.1[0-3] | 375.11 |  |
| Primary angle-closure glaucoma | H40.2 | 365.2 | 375.00\|375.11 |  |  |
| Glaucoma secondary to eye trauma | H40.3 | 365.6E |  |  |  |
| Glaucoma secondary to eye inflammation | H40.4 | 365.6F\|365.6B | 364.01 |  |  |
| Glaucoma secondary to other eye disorders | H40.5 | 365.5\|365.6A |  |  |  |
| Glaucoma secondary to drugs | H40.6 | 365.3A |  |  |  |
| Other and unspecified glaucoma | H40.[8-9] | 365.6X\|365.6C\|365.[8-9] | 375.2\|375.9 | 375.97 |  |
| Glaucoma in diseases classified elsewhere | H42 | 365.4A |  |  |  |
| Use of antiglaucoma preparations and miotics |  |  |  |  | S01EA |

**Supplementary Figure S5:**
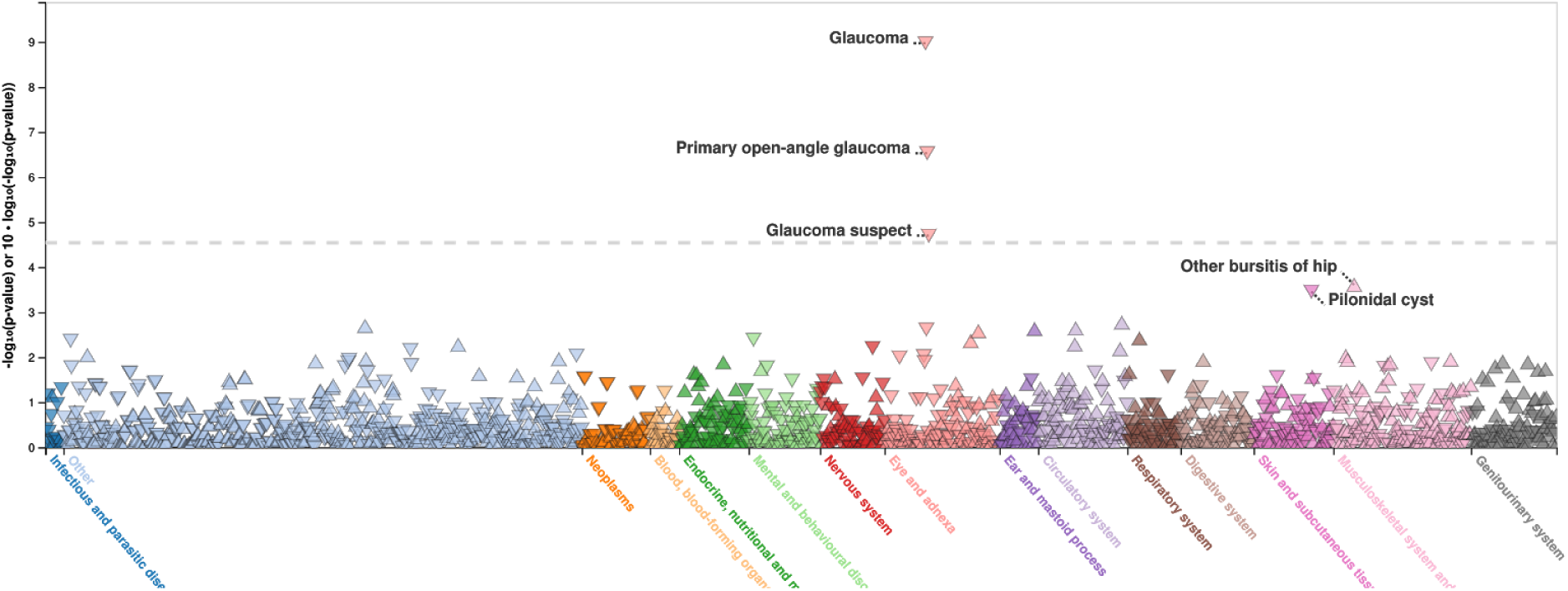
Phenome-wide association analysis of p.R220C in FinnGen. −log10(P-value) is displayed in y-axis. Disease endpoints grouped by disease categories are displayed on the x-axis. Highlighted associations with *P* < 0.001 are shown.

**Supplementary Table S6.**
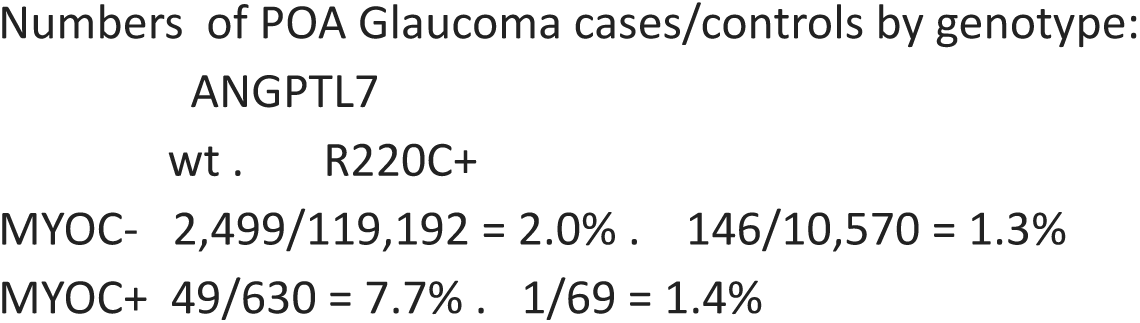
FinnGen summary of *MYOC* p.Gln168Ter and *ANGPTL7* p.Arg220Cys carriers in primary open glaucoma cases.

**Supplementary Table S7.** *ANGPTL7* protein-altering variant association with reproductive fitness, (A) the number of live births and (B) the number of children fathered. Variant includes chromosome, position, reference, and alternate allele (hg19). rsID - the rs identifier of the genetic variant. HGVSp - the HGVS protein sequence name. MAF - the minor allele frequency in UK Biobank British population. Beta - estimated regression coefficient with 95% confidence intervals. P - p-value of association.

A. Number of live births
| Variant | rsID | HGVS | MAF | Dosage | Beta [95% CI] | P |
| --- | --- | --- | --- | --- | --- | --- |
| 1:11252357:A:G | rs200058074 | p.Gln136Arg | .054 % | Gln/Arg | 0.089 [-0.064, 0.24] | 0.26 |
| 1:11252369:G:A | rs28991002 | p.Arg140His | .25 % | Arg/His | 0.29 [-1.9, 2.5] | 0.80 |
| 1:11253684:G:T | rs28991009 | p.Gln175His | .81 % | Gln/His | -0.0076 [-0.050, 0.035] | 0.72 |
|  |  |  |  | His/His | 0.23 [-0.48, 0.93] | 0.52 |
| 1:11253688:C:T | rs143435072 | p.Arg177Ter | .041 % | Arg/Ter | 0.2 [0.021, 0.38] | 0.029 |

B. Number of children fathered
| Variant | rsID | HGVS | MAF | Dosage | Beta [95% CI] | P |
| --- | --- | --- | --- | --- | --- | --- |
| 1:11252357:A:G | rs200058074 | p.Gln136Arg | .054 % | Gln/Arg | 0.0064 [-0.19, 0.21] | 0.95 |
| 1:11252369:G:A | rs28991002 | p.Arg140His | .25 % | Arg/His | 0.0030 [-0.081, 0.087] | 0.94 |
| 1:11253684:G:T | rs28991009 | p.Gln175His | .81 % | Gln/His | -0.0052 [-0.054, 0.044] | 0.83 |
|  |  |  |  | His/His | 0.41 [-0.17, 0.98] | 0.16 |
| 1:11253688:C:T | rs143435072 | p.Arg177Ter | .041 % | Arg/Ter | -0.052 [-0.27, 0.17] | 0.64 |

**Supplementary Figure S6:**
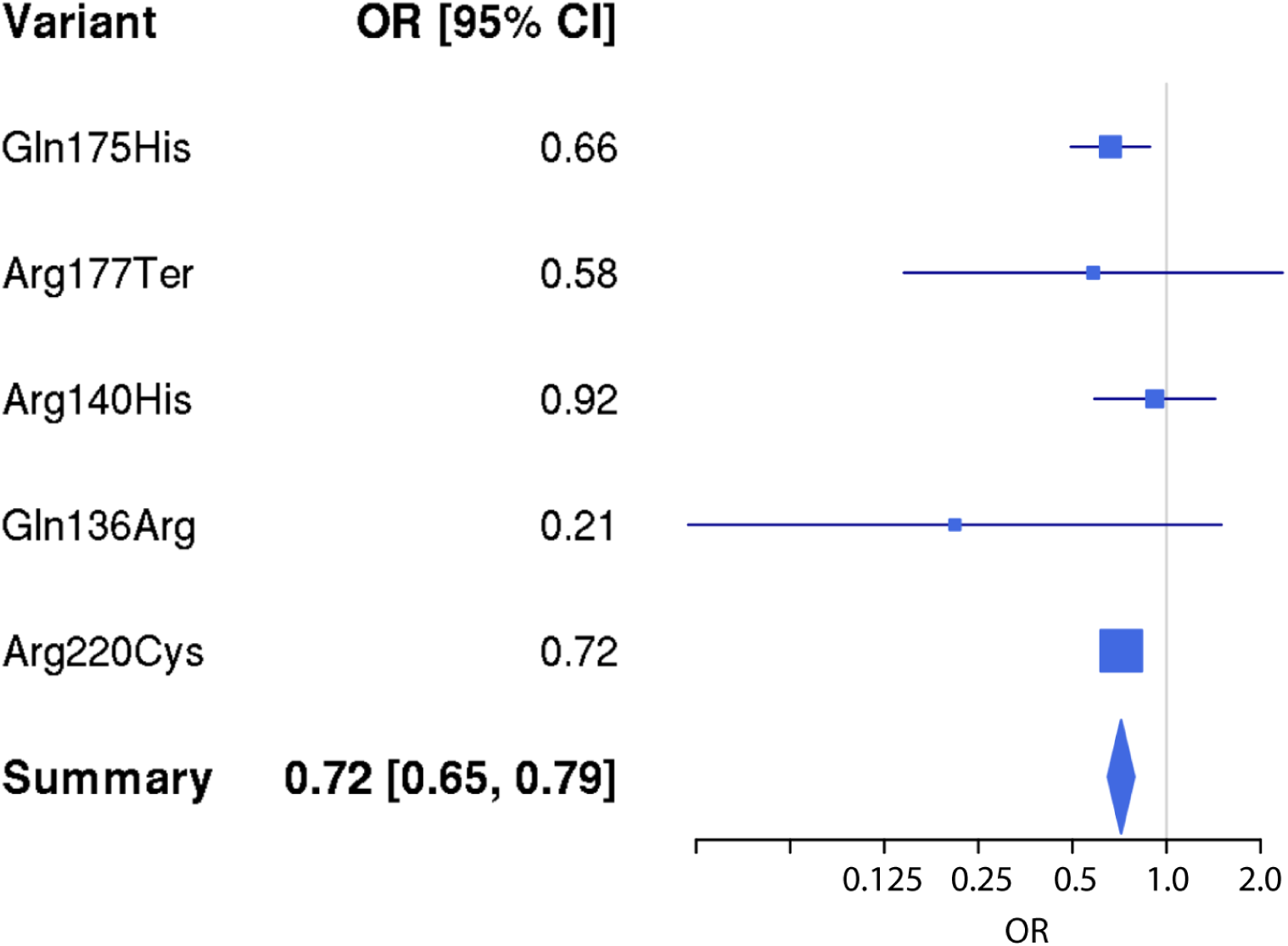
The combined effect size estimates of rare protein-altering variants. The estimates of effect size (odds ratio, OR) on glaucoma is shown in log scale for the five protein-altering variants in *ANGPTL7*. The combined effect size estimate is summarized using inverse-variance method under the fixed effect model and 95% confidence intervals are shown (Methods).

**Supplementary Figure S7:**
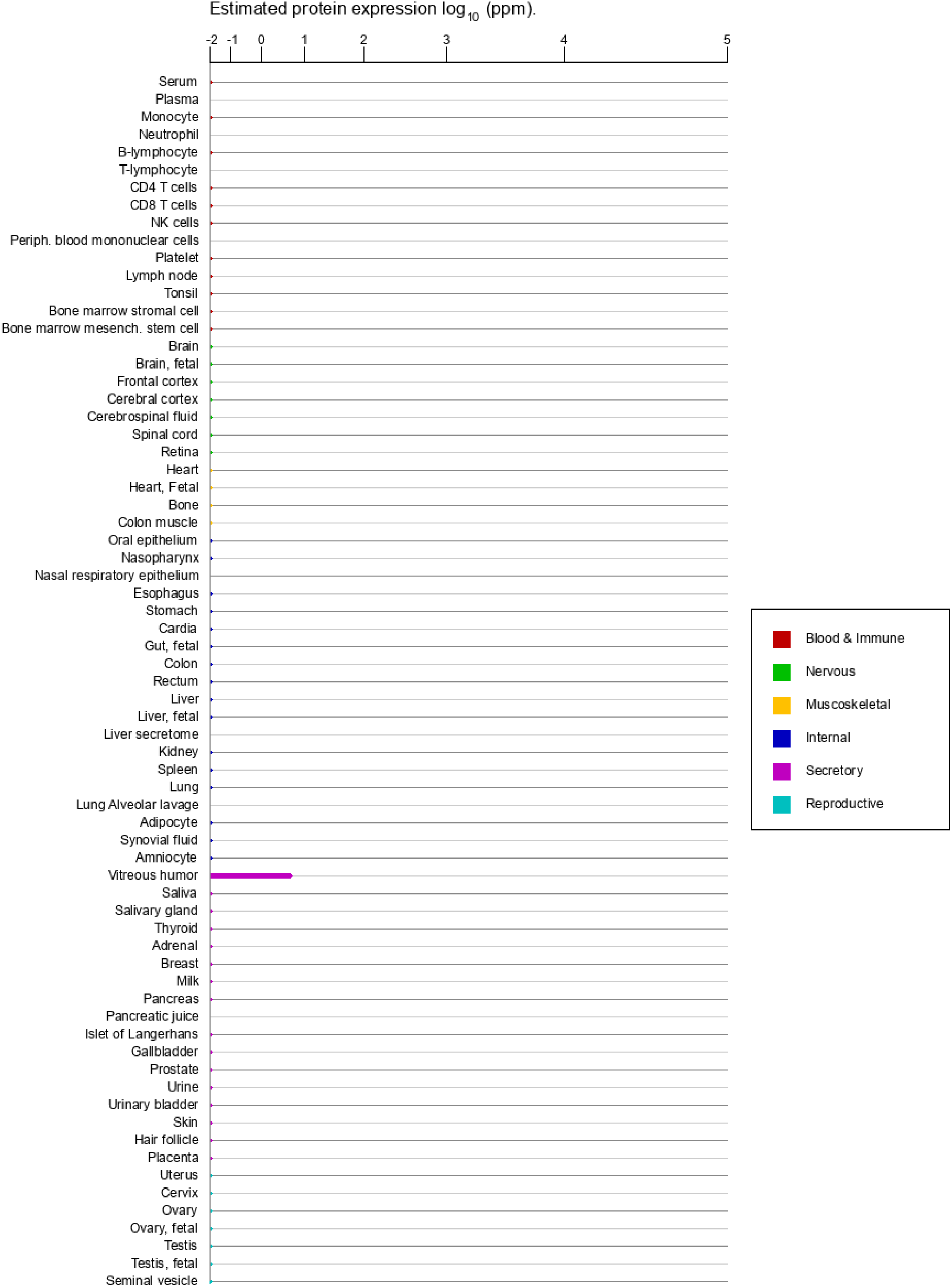
Protein expression in normal tissues and cell lines from ProteomicsDB and MOPED for ANGPTL7

## Supplementary Note

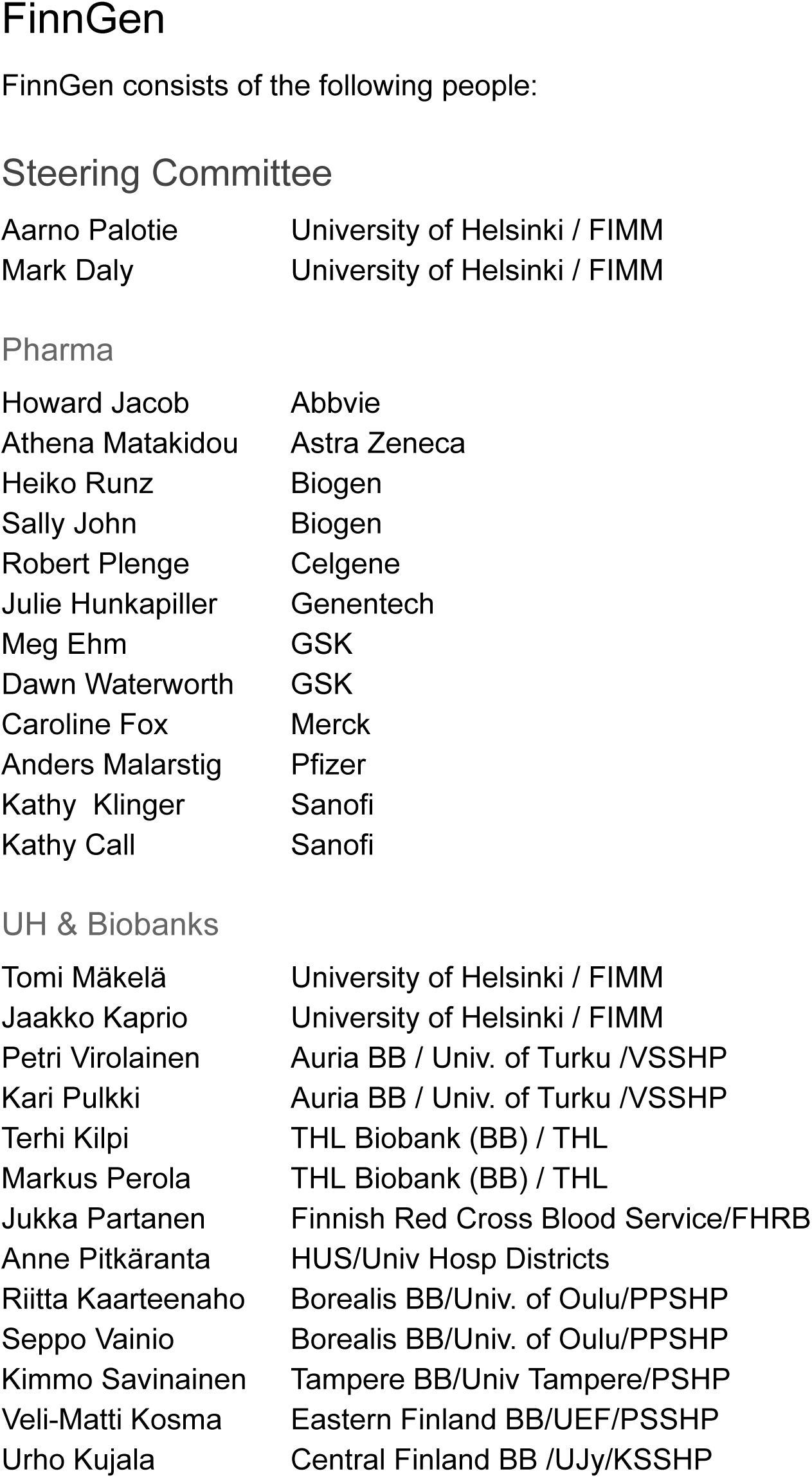
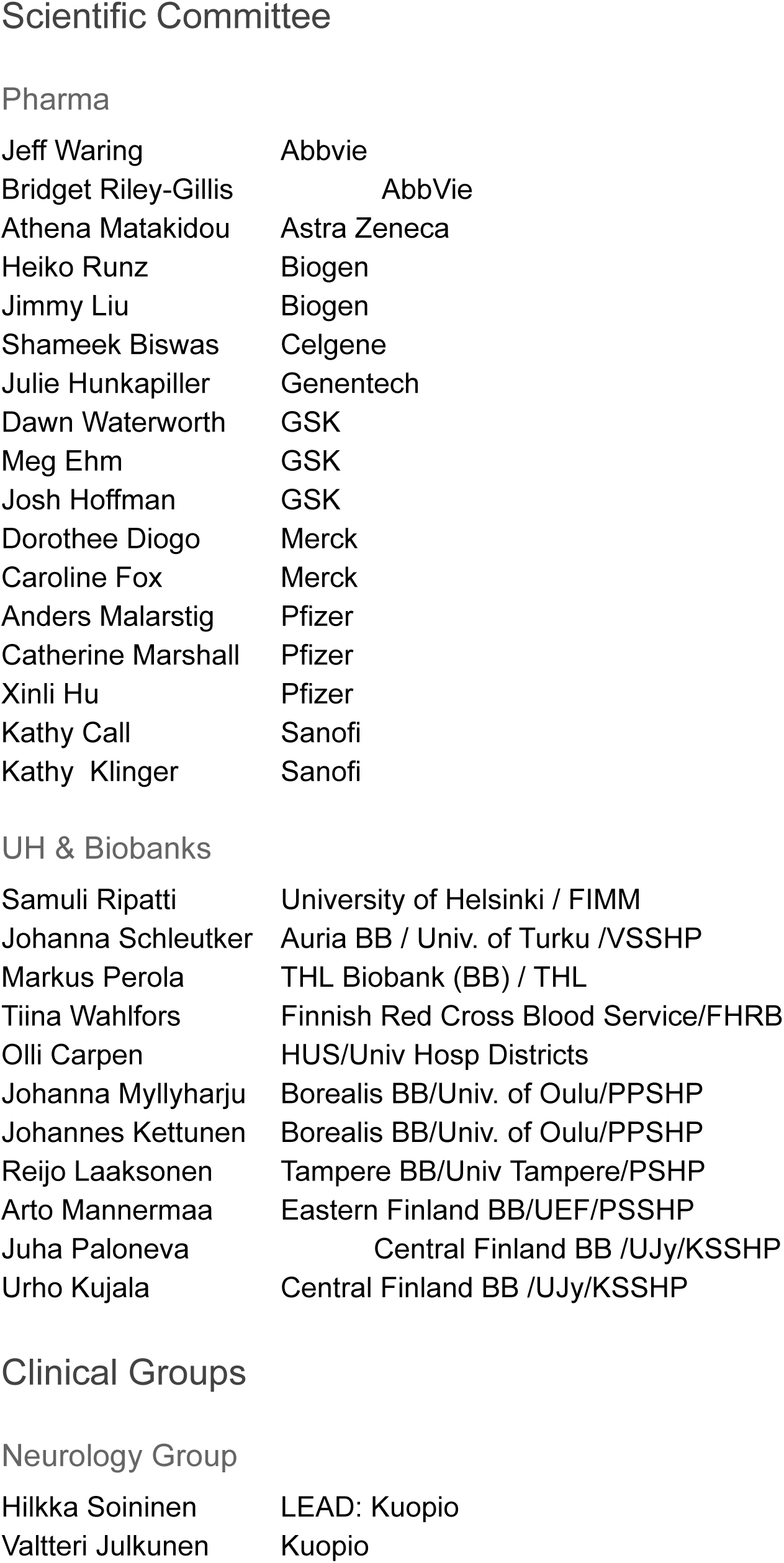
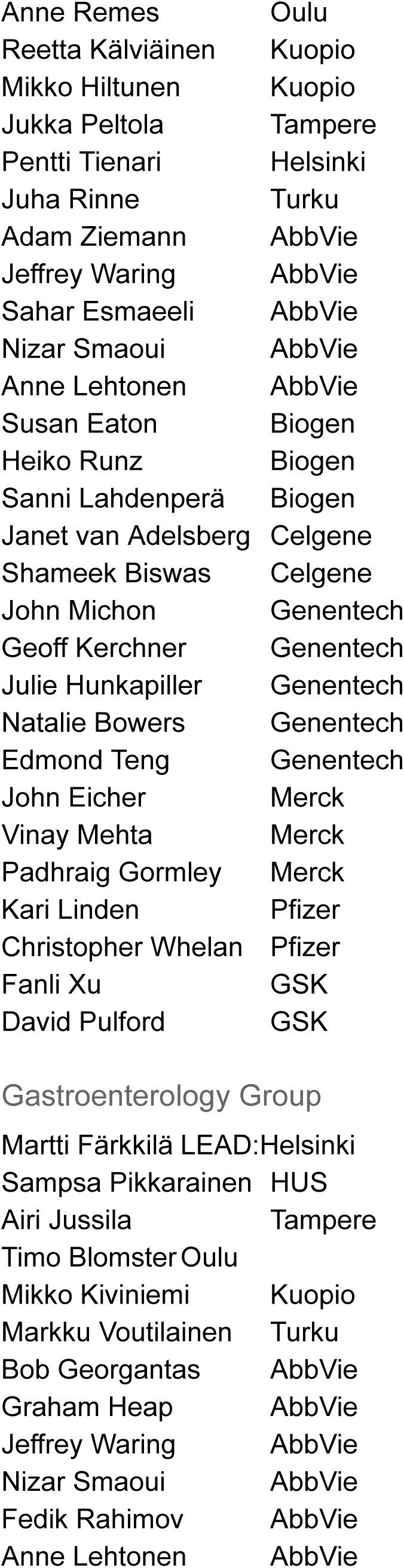
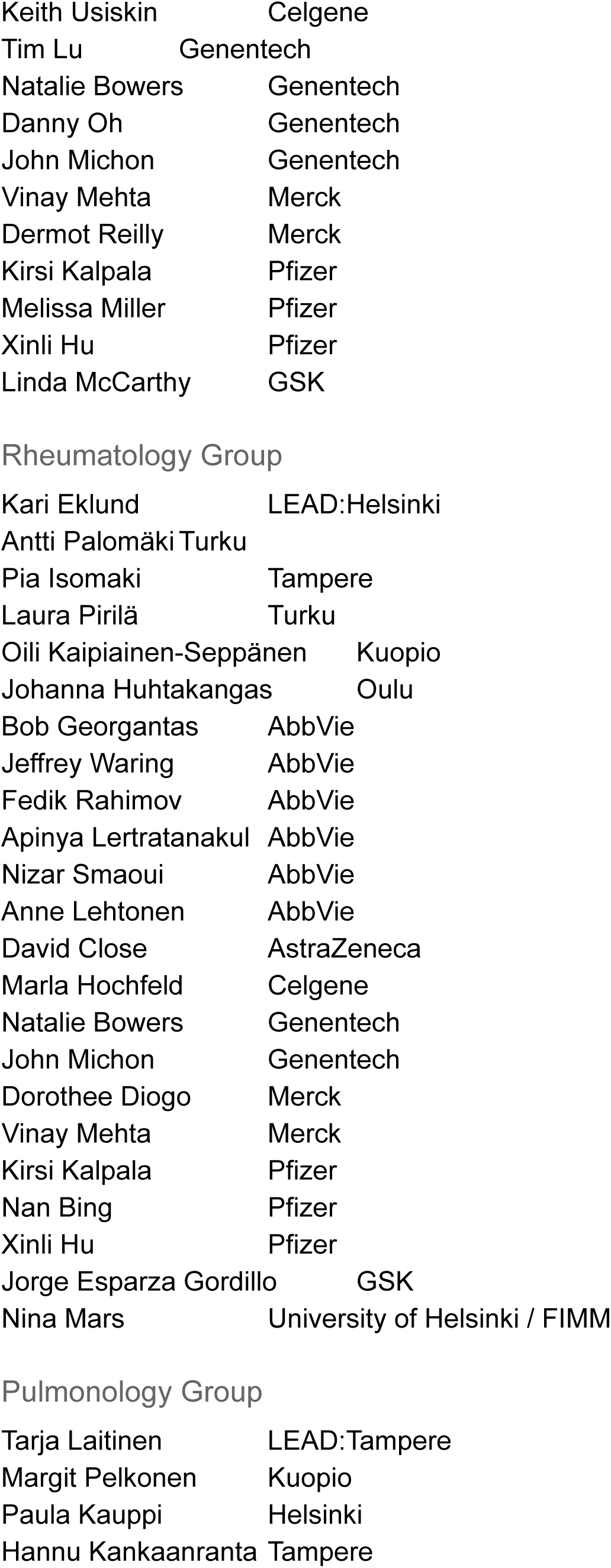
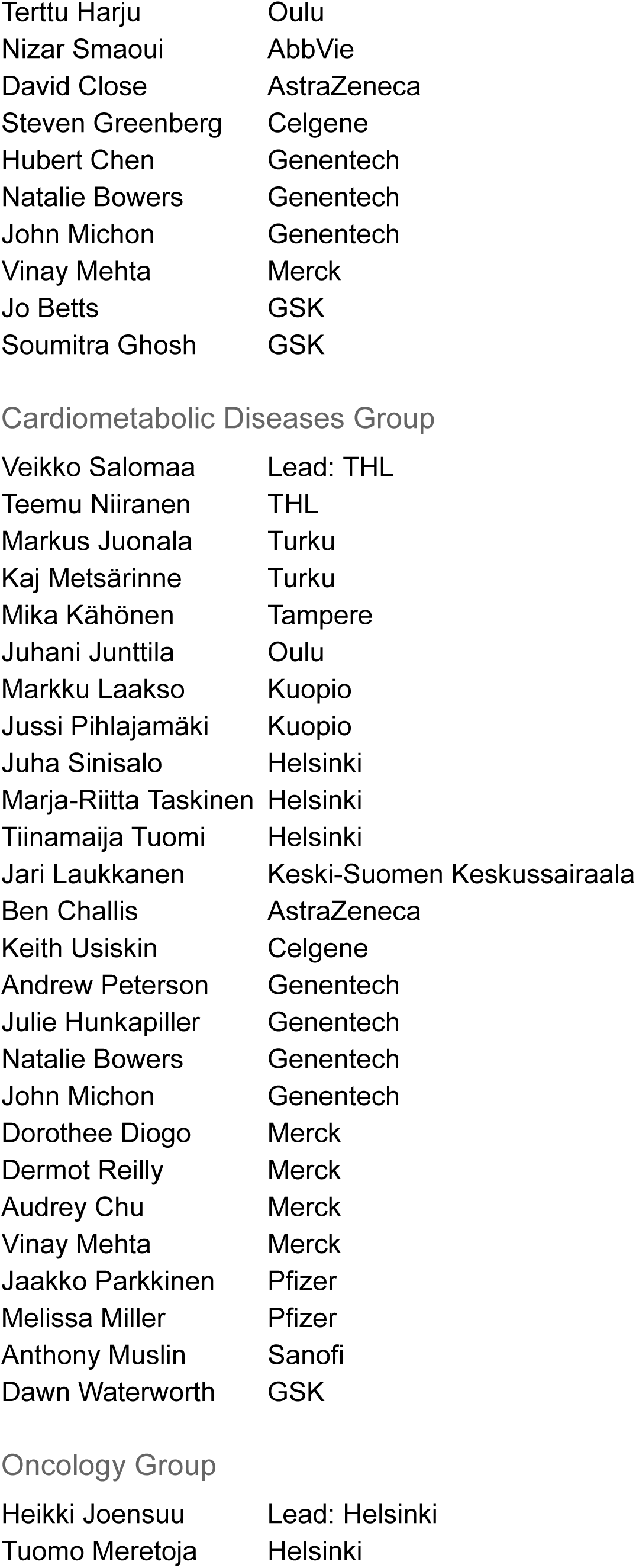
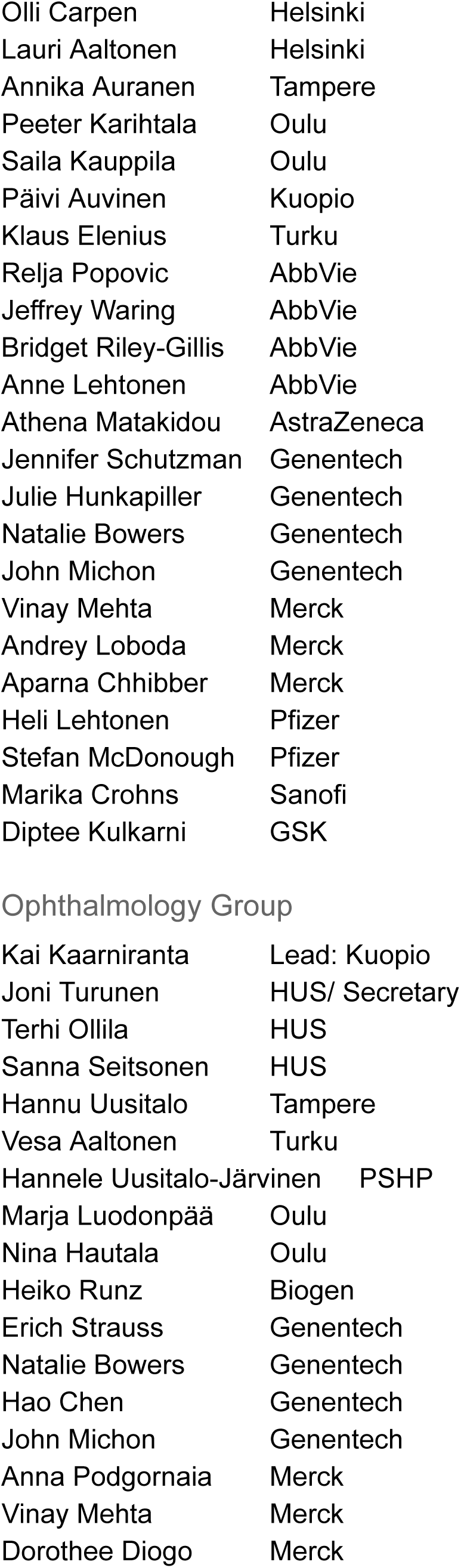
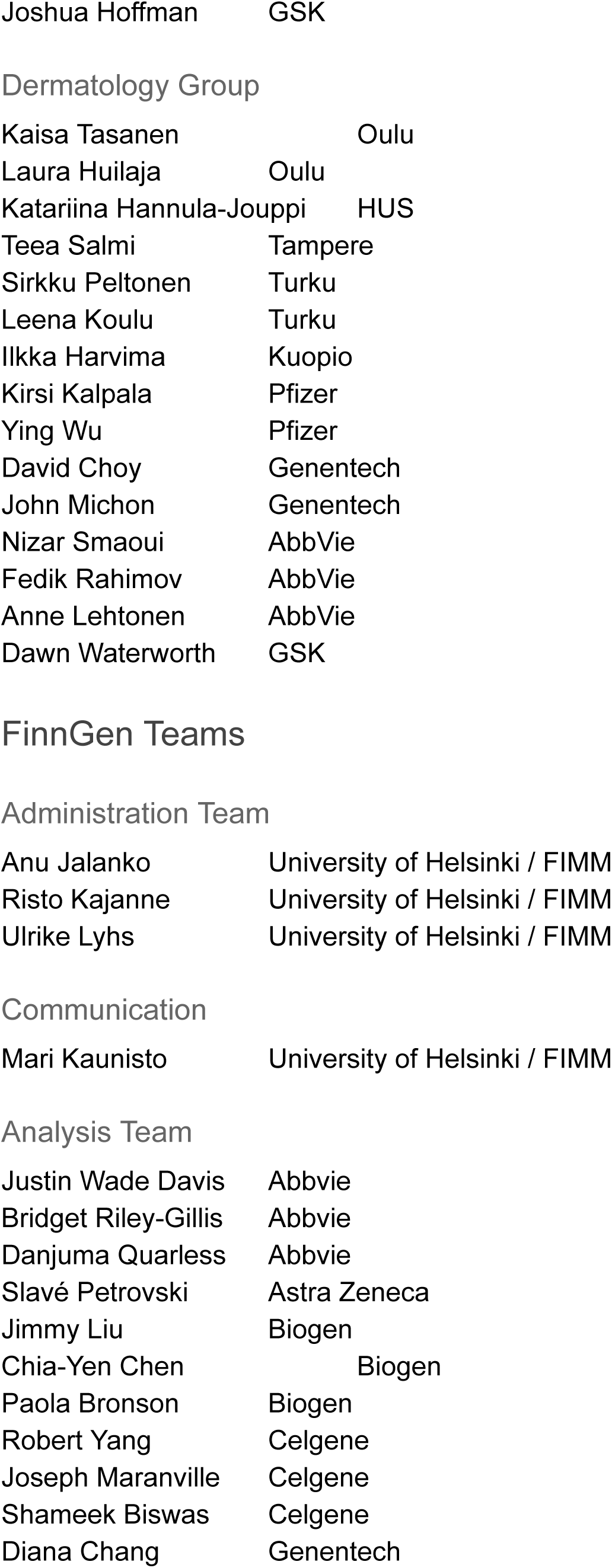
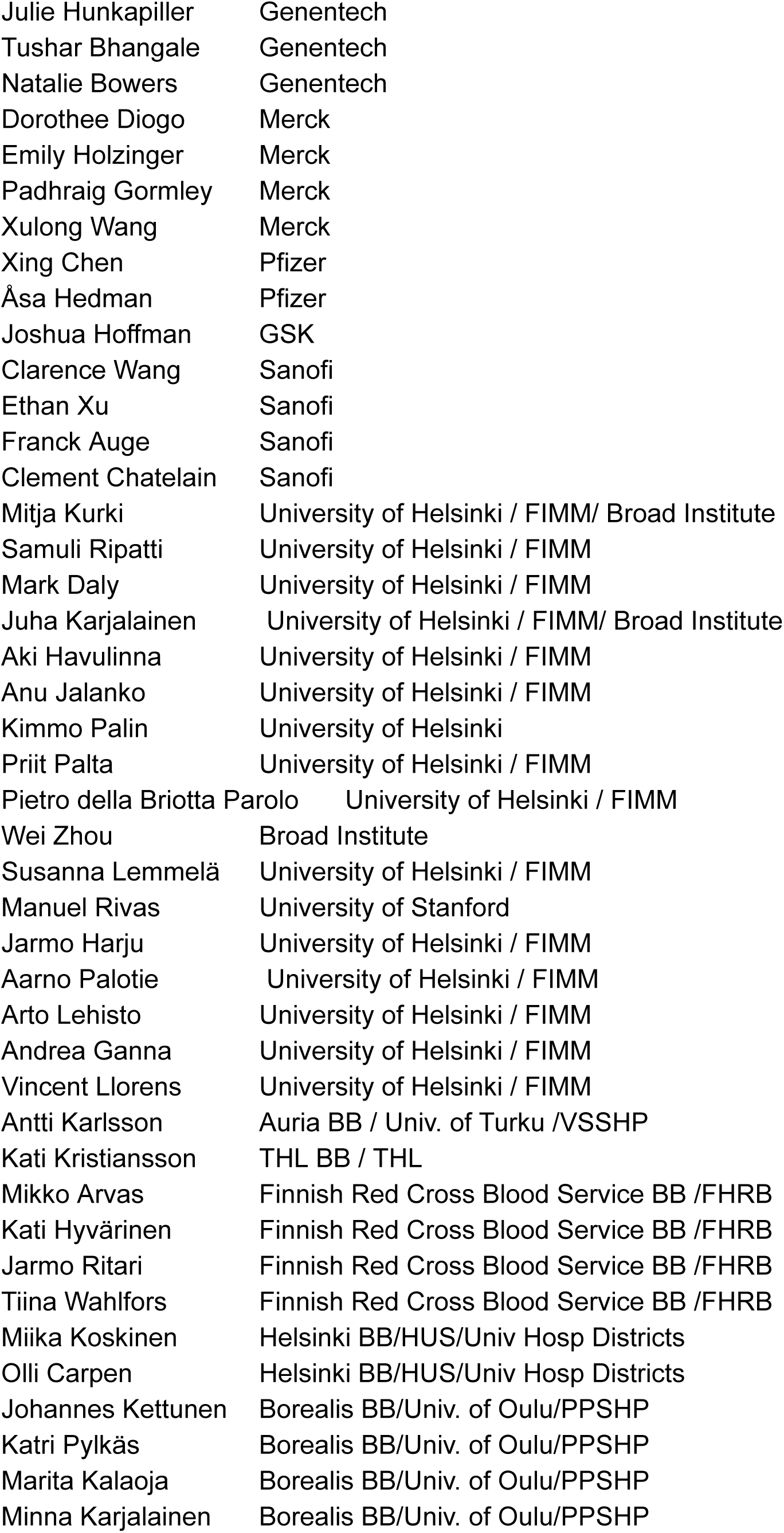
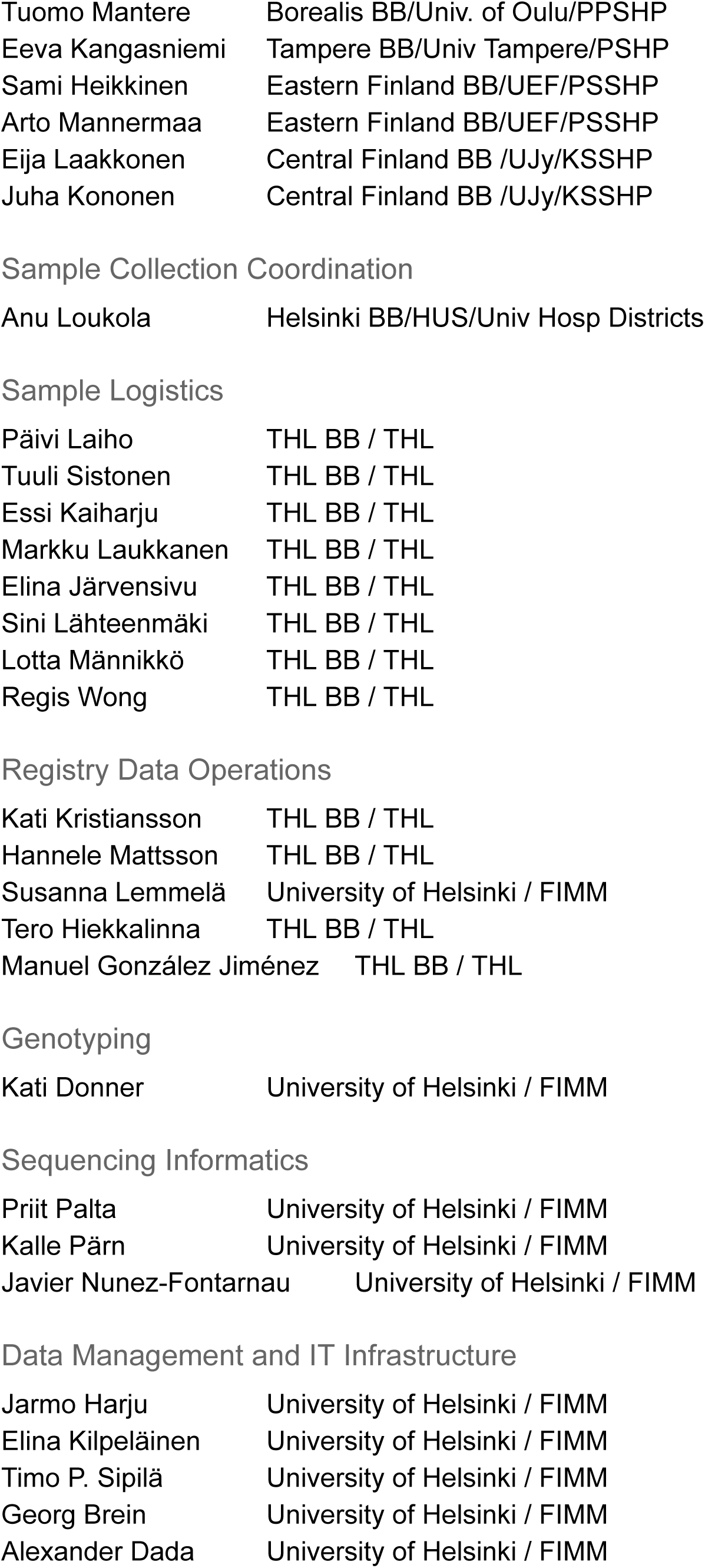
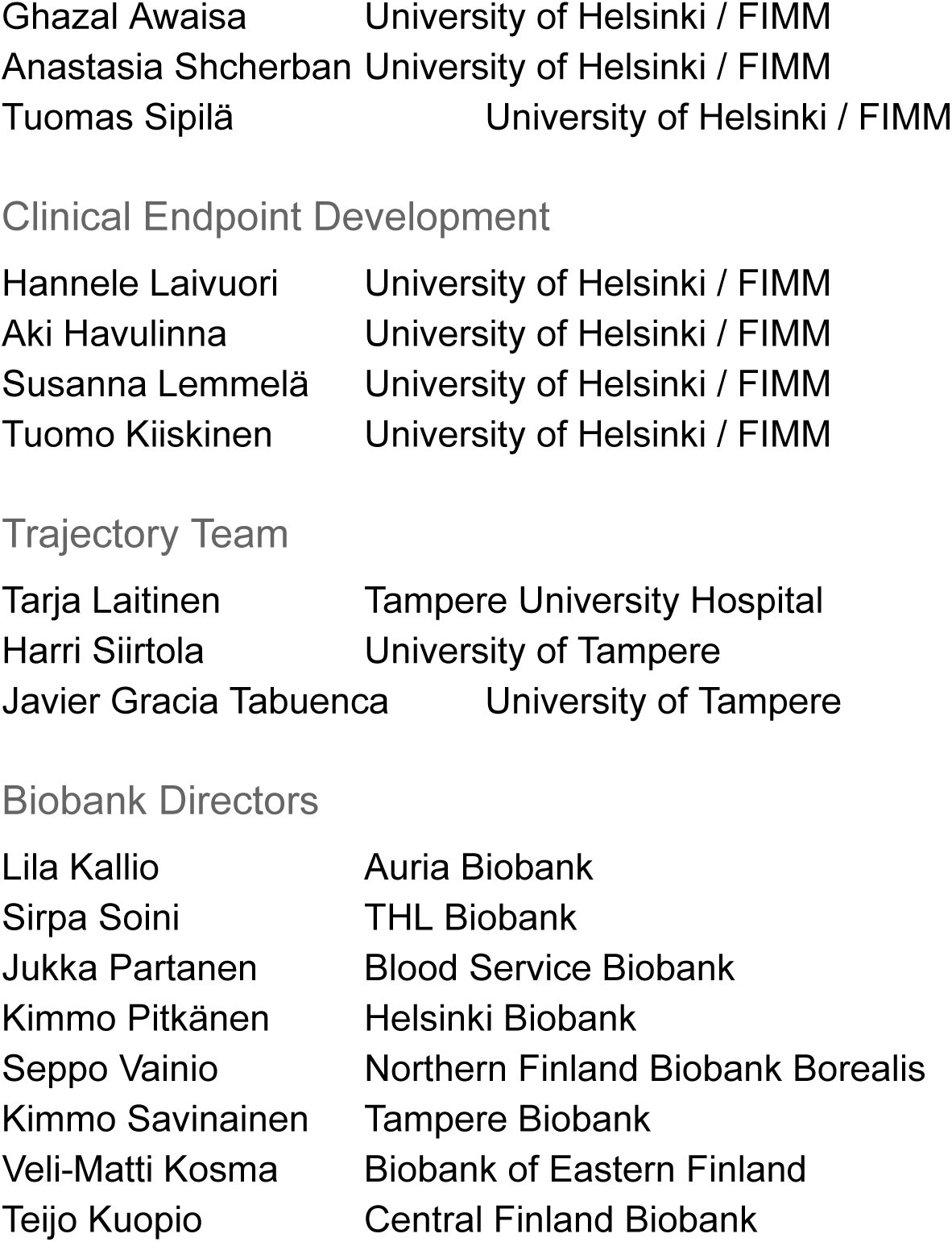

